# Mineralized collagen scaffolds fabricated with amniotic membrane matrix increase osteogenesis under inflammatory conditions

**DOI:** 10.1101/2020.01.23.917443

**Authors:** Marley J. Dewey, Eileen M. Johnson, Simona T. Slater, Derek J. Milner, Matthew B. Wheeler, Brendan A.C. Harley

## Abstract

Defects in craniofacial bones occur congenitally, after high-energy impacts, and during the course of treatment for stroke and cancer. These injuries are difficult to heal due to the overwhelming size of the injury area and the inflammatory environment surrounding the injury. Significant inflammatory response after injury may greatly inhibit regenerative healing. We have developed mineralized collagen scaffolds that can induce osteogenic differentiation and matrix biosynthesis in the absence of osteogenic media or supplemental proteins. The amniotic membrane is derived from placentas and has been recently investigated as an extracellular matrix to prevent chronic inflammation. Herein, we hypothesized that a mineralized collagen-amnion composite scaffold could increase osteogenic activity in the presence of inflammatory cytokines. We report mechanical properties of a mineralized collagen-amnion scaffold and investigated osteogenic differentiation and mineral deposition of porcine adipose derived stem cells within these scaffolds as a function of inflammatory challenge. Incorporation of amniotic membrane matrix promotes osteogenesis similarly to un-modified mineralized collagen scaffolds, and increases in mineralized collagen-amnion scaffolds under inflammatory challenge. Together, these findings suggest that a mineralized collagen-amnion scaffold may provide a beneficial environment to aid craniomaxillofacial bone repair, especially in the course of defects presenting significant inflammatory complications.

## 1. Introduction

The immune system plays a vital role in the outcome of tissue regeneration in bone defects, contributing to successful healing of the injury or inhibiting bone formation. Many strategies to promote bone regeneration exist, most commonly through the use of autografts and allografts. However, these strategies involve extensive processing of the bone substitute and residual material can be difficult to completely remove, causing an immune response (1,2). The impact of the immune system on overall outcome of implanted materials is of paramount importance: chronic and persistent inflammation can lead to fibrous tissue, limited bone formation, and possible resorption of the bone (3,4). Upon implantation of biomaterials, macrophages may encounter difficulties degrading the implant material which can lead to a foreign body reaction and ultimate failure of the implant (5). Successful healing of craniomaxillofacial (CMF) defects, specific bone defects consisting of large missing segments of bone from the head and jaw, depends greatly on the immune system behavior due to these implant-related immune responses. It is therefore desirable to develop biomaterials that can not only repair the missing bone in these defects but interact with the immune system to counteract inhibitory conditions and accelerate repair.

Mineralized collagen scaffolds have been extensively developed over the past few years to aid in the repair of CMF defects (6,7,16–21,8–15). These scaffolds are synthesized by lyophilization and offer an open pore network with the optimal pore size for cell infiltration and attachment (13,19–25). The addition of mineral to collagen scaffolds has been shown to promote osteogenesis and mineral formation *in vitro* and *in vivo*, even in the absence of osteogenic media and BMP-2 supplements to enhance bone formation (15,26). Recently, the amniotic membrane derived from placentas has been investigated as a natural extracellular matrix that could be used in biomaterial implants for its anti-inflammatory properties (27–31). In addition, amniotic membrane extracts have been shown to provide growth factors for osteogenic differentiation and repair bone defects (32–36). This membrane was investigated in non-mineralized collagen scaffolds for tendon repair due to its potential to modulate the immune response (37,38). These scaffolds demonstrated increased metabolic activity to a pro-inflammatory challenge compared to scaffolds without amniotic membrane supplementation and had a decrease in the pro-inflammatory gene, *TNF-α* (37). Although the amniotic membrane was successfully incorporated into non-mineralized collagen scaffolds for tendon repair, scaffolds lacking mineral components have been shown to be less than ideal for bone regeneration (26). As a result, it may be beneficial to incorporate amniotic membrane matrix within mineralized collagen scaffolds to achieve both the osteogenic and immunomodulatory capabilities we desire in a biomaterial for bone repair.

In this manuscript, we report a mineralized collagen-amnion scaffold that can promote osteogenesis, even under inflammatory challenge. We hypothesized that the addition of the amniotic membrane to mineralized collagen scaffolds would increase osteogenesis and in addition, it would continue to promote osteogenesis in *in vitro* culture containing inflammatory cytokines. We describe the mechanical properties and pore structure of mineralized collagen scaffolds containing the amniotic membrane. Subsequently, we evaluate the *in vitro* behavior of porcine adipose derived stem cells (pASC) seeded on these scaffolds in normal growth medium and medium supplemented with an inflammatory protein.

## 2. Materials and Methods

### 2.1. Experimental design

In order to test our first hypothesis that mineralized collagen scaffolds containing the amniotic membrane would promote osteogenesis, porcine adipose derived stem cells (pASC) were seeded onto mineralized collagen-amnion scaffolds and mineralized collagen scaffolds. Cell viability, osteogenic gene expression, osteogenic protein activity, and mineral content were evaluated over 28 days. To test our second hypothesis that scaffolds containing the amnion would continue to promote osteogenesis in inflammatory conditions, mineralized collagen-amnion scaffolds were placed in normal growth medium and inflammatory medium. Cell viability, osteogenic gene expression, osteogenic protein activity, and mineral content were evaluated over 28 days (**Fig. 1**).

**Figure 1.**
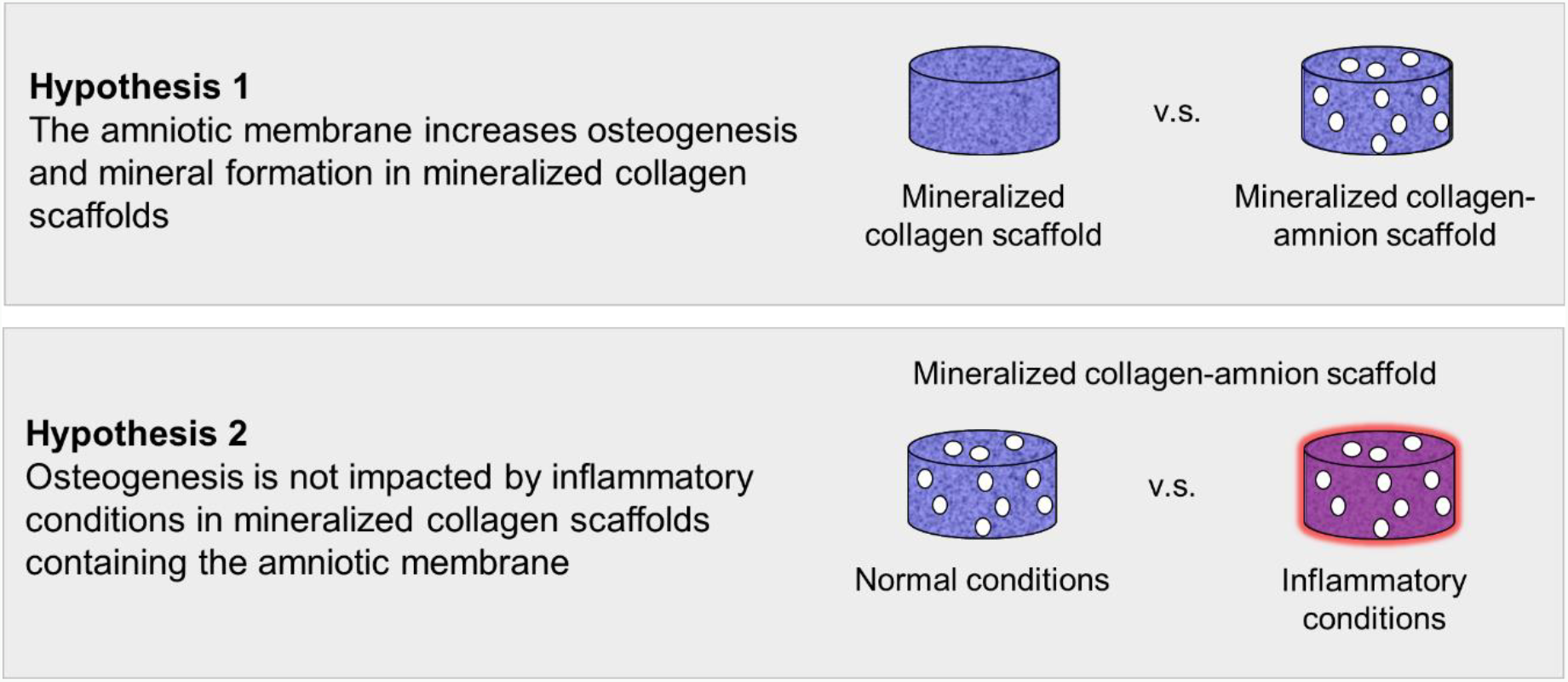
Experimental outline. Hypothesis 1: mineralized collagen-amnion scaffolds will further promote osteogenesis and mineral formation compared to mineralized collagen scaffolds. Hypothesis 2: mineralized collagen-amnion scaffolds placed in inflammatory conditions (normal growth medium supplemented with IL-1β) will continue to promote osteogenesis due to the immunomodulatory nature of the amniotic membrane.

### 2.2. Isolation of the amniotic membrane from human placentas

Human placentas were obtained from a collaboration with Carle Foundation Hospital Tissue Repository (Urbana, IL) once having met set standards of uncomplicated vaginal births. The amniotic membrane was isolated from placentas as previously described (37,38). Briefly, the amniotic membrane was separated mechanically from the placenta and blood on the membrane was washed away in calcium- and magnesium-free HBSS. The amnion was then scraped with a spatula to remove any remaining blood and parts of the spongy layer. The amnion was subsequently cut into smaller sections and decellularized in 125 μg/mL thermolysin (Sigma-Aldrich, St. Louis, MO) (39). The decellularized amniotic membrane pieces were rinsed in PBS to remove any debris and were stored in PBS at 4°C for 24-48 hours to allow the amnion to swell. After soaking in PBS, the amnion was once again scraped with a spatula to remove the spongy layer, and then the amniotic membrane segments were lyophilized and stored in a desiccator until use (**Fig. 2**).

**Figure 2.**
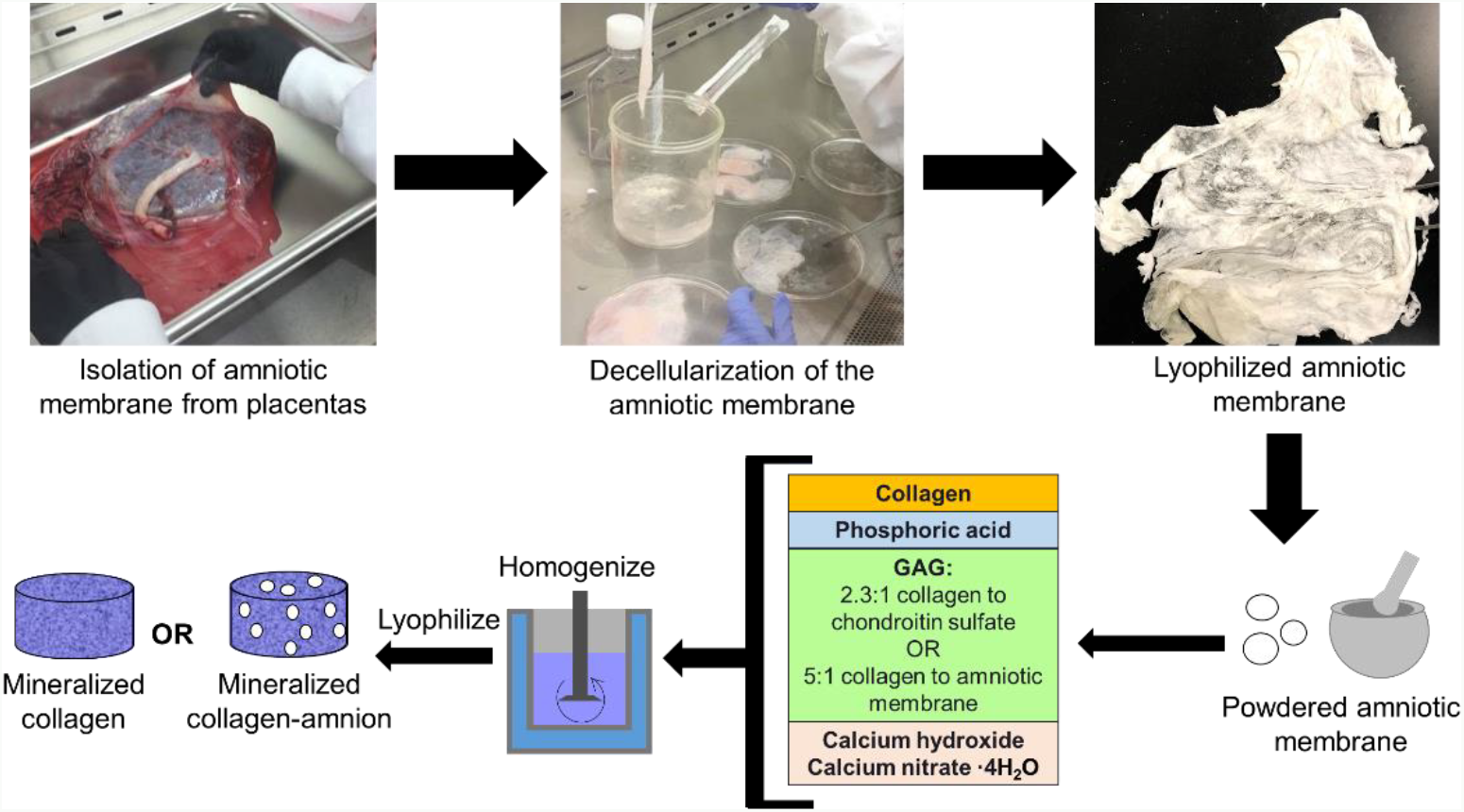
Isolation of amniotic membrane from placentas and synthesis of mineralized collagen and mineralized collagen-amnion scaffolds.

### 2.3. Fabrication of mineralized collagen and mineralized collagen-amnion scaffolds

#### 2.3.1. Fabrication of mineralized collagen scaffolds

Mineralized collagen scaffolds were fabricated by lyophilizing a mineralized collagen suspension as previously described (40). The suspension, made of 1.9 w/v% bovine type I collagen (Collagen Matrix, Oakland, NJ), 0.84 w/v% chondroitin-6-sulfate (Sigma-Aldrich), and calcium salts (Ca(OH)2, Ca(NO3)2·4H2O, Sigma-Aldrich) in phosphoric acid (Sigma-Aldrich), was homogenized then stored at 4°C. Prior to use, the suspension was degassed then lyophilized via a Genesis freeze-dryer (VirTis, Gardener, NY) (21). For *in vitro* testing, approximately 24 mL of suspension was pipetted into an aluminum square mold (76.2 mm width and 19.05 mm height). For mechanical testing, approximately 1 mL of slurry was pipetted into polysulfone molds measuring 10 mm high and 11.9 mm wide. Lyophilization proceeded by freezing at a constant temperature decrease of 1°C/min starting from 20°C and ending at −10°C to form ice crystals. Then the suspension was held at −10°C for 2 hours and sublimated by decreasing the pressure to 0.2 Torr and 0°C, producing a porous scaffold network by evaporating the ice crystals. Finally, the scaffolds were allowed to come to room temperature and atmospheric pressure. After lyophilization, the sheet of mineralized collagen was removed from the plate and scaffolds were punched out of the sheet using a 6 mm biopsy punch (Integra, Plainsboro, NJ) and were cut to 3 mm heights with a razor blade if necessary.

#### 2.3.2. Fabrication of mineralized collagen-amnion scaffolds

Mineralized collagen-amnion scaffolds were fabricated in a similar manner to mineralized collagen scaffolds as described above. Before adding the amniotic membrane to mineralized collagen-amnion scaffolds, it was first ground with a mortar and pestle with a small amount of phosphoric acid and calcium hydroxide solution to create fine particles of matrix before being introduced into the suspension. The mineralized collagen-amnion suspension was made of 1.9 w/v% bovine type I collagen (Collagen Matrix), a 5:1 w:w ratio of collagen:amnion due to the high collagen content in the amniotic membrane (37), and calcium salts (Ca(OH)2, Ca(NO3)2·4H2O, Sigma-Aldrich) in phosphoric acid (Sigma-Aldrich). After homogenizing, it was lyophilized following the same procedure as mineralized collagen scaffolds, however, 40-50 ml of slurry was used in the square aluminum mold to cover the same area and depth as mineralized collagen scaffolds (**Fig. 2**).

### 2.4. SEM imaging

An environmental scanning electron microscope (SEM) was utilized to visualize the pore architecture of mineralized collagen and mineralized collagen-amnion scaffolds without cells. Scaffolds were cut in half using a razor blade, exposing the interior, then were sputter coated with Au/Pd prior to visualizing using a FEI Quanta FEG 450 ESEM (FEI, Hillsboro, OR).

### 2.5. Pore size analysis

The pore size of mineralized collagen and mineralized collagen-amnion scaffolds was analyzed following a JB-4 (Polysciences, Inc., Warrington, PA) embedding procedure (9,25). Briefly, scaffolds were hydrated in 100% ethanol under vacuum inside a desiccator before embedding. After soaking in JB-4, the scaffolds were placed into wells to harden, with three samples of each type placed flat into the mold and three placed on their side to create transverse and longitudinal sections. These molds were then placed at 4°C overnight to complete polymerization. JB-4 embedded scaffolds were embedded in paraffin to fit in molds for microtome sectioning, and 5 μm sections were cut using a RM2255 microtome (Leica, Wetzlar, Germany) with a tungsten carbide blade. Scaffolds were sectioned and placed onto glass slides, then stained with an aniline blue solution (Thermo Fisher Scientific, Waltham, MA). Slides were then imaged with a NanoZoomer Digital Pathology System (Hamamatsu, Japan). To analyze pore structure, images were captured of each section and images were analyzed by a custom Matlab pore size code (9,41) to get an average pore size and aspect ratio for each scaffold.

### 2.6. Mechanical compression testing of mineralized collagen and mineralized collagen-amnion scaffolds

The mechanical compressive behavior of the mineralized collagen and mineralized collagen-amnion scaffolds without cells was analyzed using an Instron 5943 mechanical tester (Instron, Norwood, MA) with a 100 N load cell under dry conditions to generate stress-strain curves for the scaffolds. Briefly, 8 samples of mineralized collagen and mineralized collagen-amnion scaffolds were compressed to failure (2 mm/min) and the collapsed stress, strain, and Young’s Modulus were determined from resulting curves by using low-density open-cell foam analysis (12,19,42). Previously, it has been shown that the hydration and EDAC crosslinking of mineralized collagen scaffolds lowers the Young’s Modulus and compressive stress (12, 19); it can be assumed that the scaffolds presented here would exhibit similar behavior.

### 2.7. Hydration, crosslinking, and sterilization of mineralized collagen and mineralized collagen-amnion scaffolds

Mineralized collagen and mineralized collagen-amnion scaffolds were sterilized using an ethylene oxide treatment for 12 hours via an AN74i Anprolene gas sterilizer (Andersen Sterilizers Inc., Haw River, NC) in sterilization pouches. All handling steps were done utilizing sterile techniques. Before seeding with cells, hydration followed previously described methods (43). Briefly, scaffolds underwent a soaking period in 100% ethanol, then multiple washes in PBS and crosslinked with a carbodiimide chemistry. Afterwards, scaffolds were washed in PBS and soaked in normal growth media for 42 hours before cell seeding.

### 2.8. Porcine adipose derived stem cell culture (pASC) and seeding on scaffolds in normal and inflammatory media

Mineralized collagen and mineralized collagen-amnion scaffolds were compared *in vitro* in a two-part study described in the experimental design section. Porcine adipose derived stem cells were expanded at 37°C and 5% CO_2_ in normal mesenchymal stem cell growth media (low glucose DMEM, 10% mesenchymal stem cell fetal bovine serum (Gibco, Waltham, MA), and 1% antibiotic-antimycotic (Gibco)) without osteogenic supplements. pASC have been shown to promote osteogenesis on mineralized collagen scaffolds in the absence of osteogenic supplements (41). pASC were used at passage 6 and each scaffold was seeded on one side with 5,000 cells/μL for 30 min, then 5,000 cells/μL on the opposite side of the scaffold for 1.5 hours, for a total of 100,000 cells per scaffold. After the initial attachment period was finished, additional normal growth media was added to each well. This marked day (−1) of the study, following previously described methods of culturing with pro-inflammatory media (37). The next day, medium was removed and the scaffolds designated for study in normal growth media were placed in wells with fresh normal growth medium and the mineralized collagen-AM scaffolds designated for study in inflammatory conditions were placed in wells with inflammatory medium (1 ng/ml of IL-1β recombinant porcine protein (R&D Systems, Minneapolis, MN) added to normal growth media), which marked day 0 of the study. During the inflammatory phase of wound healing, the pro-inflammatory cytokine IL-1β is secrete by macrophages and represents inflammatory stimuli in this study. The study then proceeded for 28 days with appropriate media replacement every third day.

### 2.9. Metabolic activity measurement of pASC on scaffolds

Metabolic activity of pASC seeded on mineralized collagen and mineralized collagen-amnion scaffolds was calculated using a non-destructive alamarBlue® assay over the course of 28 days (day 0, 4, 7, 14, 28). Six scaffolds were rinsed in PBS prior to incubation under gentle shaking in alamarBlue® (Invitrogen, Carlsbad, California) in an incubator at 37°C for 2 hours. Following incubation, the alamarBlue® solution was measured using a F200 spectrophotometer (Tecan, Mannedorf, Switzerland) for the fluorescence of resorufin (540(52) nm excitation, 580(20) nm emission). Scaffold metabolic activity at days 0 through 28 was calculated from a standard curve generated on day (−1) of known numbers of cells and normalized to the cell seeding density of 100,000 cells.

### 2.10. Cell number of pASC on scaffolds

Total cell number of pASC on mineralized collagen and mineralized collagen-amnion scaffolds was calculated using a Hoechst 33258 (Invitrogen) DNA quantification method over the course of 28 days (day 0, 7, 14, 28) (44). Five seeded scaffolds of each scaffold type were rinsed in PBS to remove any unattached cells, and then placed in a papain solution (Sigma-Aldrich) for 24 hours in a 60°C water bath to digest and lyse cells. Hoechst dye fluorescently labels double-stranded DNA and a spectrophotometer (Tecan) (360 nm excitation, 465 nm emission) was used to measure this fluorescence. The total number of cells in each scaffold was normalized to the fluorescence of an unseeded control scaffold at each timepoint, and the cell number was calculated from fluorescent reading by a standard generated with known cell number at day (−1) and normalized to the cell seeding density of 100,000 cells.

### 2.11. Western Blot protein activity analysis of scaffolds

Three of each cell-seeded mineralized collagen and mineralized collagen-amnion scaffold were lysed for protein at days 0, 4, 7, 14, 28. To extract protein, a solution of phosphate inhibitor cocktails (Sigma-Aldrich) and a RIPA lysis buffer were used (45). To evaluate the protein concentration, a Pierce™ BCA Protein Assay Kit (Thermo Fisher Scientific) and a Pierce™ Bovine Serum Albumin Standard Pre-Diluted Set (Thermo Fisher Scientific) were used. Analysis of protein activity was quantified with a Western Blot assay, loading 5 μg of protein lysate in each lane and using a SuperSignal West Femto Maximum Sensitivity Substrate (Thermo Fisher Scientific) or a SuperSignal West Pico PLUS Sensitivity Substrate (Thermo Fisher Scientific) to visualize bands. Images were captured with an Image Quant LAS 4010 machine (GE Healthcare Life Sciences, Little Chalfont, United Kingdom). Primary and secondary antibodies are listed in **Table 1**, and β-actin was used as a loading control.

**Table 1.** Antibodies used in Western Blots

| Protein | Blocking | Primary Antibody | Secondary Antibody |
| --- | --- | --- | --- |
| β-actin<br>(45 kDa) | 5% dry milk in TBST | 1:1000 in 5% dry milk<br>(Cell signaling<br>Technologies, Rabbit<br>mAB, 4967) | 1:2500 in TBST, Anti-Rabbit IgG, HRP<br>linked antibody (Cell<br>Signaling<br>Technologies, 7074) |
| ERK1/2<br>(42-44 kDa) |  | 1:1000 in 5% dry milk<br>(Cell signaling<br>Technologies, Rabbit<br>mAB, 9102) |  |
| p-ERK1/2<br>(44-42 kDa) |  | 1:1000 in 5% dry milk<br>(Cell signaling<br>Technologies, Rabbit<br>mAB, 9101) |  |
| p38<br>(40 kDa) |  | 1:1000 in 5% dry milk<br>(Cell signaling<br>Technologies, Rabbit<br>mAB, 9211) |  |
| p-p38<br>(43 kDa) |  | 1:1000 in 5% dry milk<br>(Cell signaling<br>Technologies, Rabbit<br>mAB, 9215) |  |
| AKT<br>(60 kDa) |  | 1:1000 in 5% dry milk<br>(Cell signaling<br>Technologies, Rabbit<br>mAB, 9272) |  |
| p-AKT<br>(60 kDa) |  | 1:2000 in 5% dry milk<br>(Cell signaling<br>Technologies, Rabbit<br>mAB, 4060) |  |

### 2.12. RT-PCR gene expression analysis of scaffolds

RNA was isolated from five mineralized collagen and mineralized collagen-amnion scaffolds across 28 days (day 0, 7, 14, 28). A RNeasy Plant Mini kit (Qiagen, Valencia, California) was used to isolate RNA, and RNA was reverse-transcribed to cDNA using a QuantiTect Reverse Transcription kit (Qiagen) and a S100 thermal cycler (Bio-Rad, Hercules, California). Samples for real-time PCR were performed in duplicate with 10 ng of cDNA with either Taqman fast advanced master mix and Taqman gene expression assays (Applied Biosystems, Foster City, California) or SSoAdvancedTM Universal SYBR® Green Supermix (Bio-Rad) and PrimePCR SYBR® Green Assay (Bio-Rad) (**Table 2**). SYBR® Green reagents were used only for the *RUNX2* gene due to limited availability as a Taqman gene expression assay. All Taqman and PrimePCR SYBR® assays were pre-validated by associated companies. PCR plates were read using a QuantstudioTM 7 Flex Real-Tim PCR System (Thermo Fisher Scientific) and results were analyzed using a delta-delta CT method. All results were expressed as fold changes, which were normalized to cell expression before seeding on scaffolds.

**Table 2.** Primers for RT-PCR

| Gene | Primer Information |
| --- | --- |
| *GAPDH* | Taqman (Ss03375629_u1)<br>Bio-Rad (PrimePCR assay, GAPDH, Ssc) |
| *COL1A2* | Taqman (Ss03375009_u1) |
| *BGLAP* | Taqman (Ss03373655_s1) |
| *BMP2* | Taqman (Ss03373798_g1) |
| Osterix (LOC404701) | Taqman (Ss03373734_s1) |
| *IL-8* | Taqman (Ss03392437_m1) |
| *RUNX2* | Bio-Rad (PrimePCR assay, RUNX2, Ssc) |

### 2.13. Micro-CT analysis of scaffolds

Mineral intensity of three mineralized collagen and mineralized collagen-amnion scaffolds was quantified at day 28 in seeded and day (−1) unseeded samples using microcomputed tomography (micro-CT). Prior to analysis, samples were fixed in 10% formalin and stored in 4°C for a minimum of 24 hours. Micro-CT was performed using a MicroXCT-400 (Zeiss, Oberkochen, Germany) and scans utilized a 1x camera, 8 W, and 40kV, with the same source and detector distance, exposure time, and binning for each scaffold. The contrast and brightness histogram measured the same value for each scaffold. ImageJ was used to evaluate the mineral content from z-stacks of 2D images generated from micro-CT by thresholding at an intensity of 241 and analyzing the particle average intensity, following a previously described procedure (46). Average fill of the scaffolds was calculated by dividing the average total area of the particles by the area of the scaffold; representative images can be seen in **Supp. Fig. 1**.

### 2.14. Histological evaluation of scaffolds

Post micro-CT analysis, three samples of each scaffold type were rinsed in PBS and then embedded in Tissue-Tek® O.C.T. compound (Sakura Finetek, Netherlands) and stored at −80°C. Slices measuring 14 μm in thickness were sectioned using a CM1900 microtome (Leica) and mounted to glass slides. Slides were then stained with Hematoxylin (Leica) and Eosin (Thermo Fisher Scientific), Alizarin Red S (Sigma-Aldrich), or Von Kossa (Abcam, Cambridge, UK). Immunohistochemistry was performed using a polyclonal antibody to osteopontin (OPN, ab8448, Abcam) and methyl green (Sigma-Aldrich) with a goat pAB to Rb IgG (HRP) (ab6721, Abcam) secondary antibody in normal goat serum (Jackson ImmunoResearch, West Grove, PA). Methyl green (Sigma- Aldrich) was used as a counterstain. All histologically stained slides were analyzed qualitatively and imaged using a NanoZoomer Digital Pathology System (Hamamatsu, Japan).

### 2.15. Statistics

Statistics were performed using OriginPro software (Northampton, Massachusetts) with significance set to p < 0.05 and tested in accordance with literature (47). A normality test was first performed using the Shapiro-Wilk test and any sample data sets that were found to be non-normal underwent a Grubbs outlier test to remove any outliers and normality was re-assessed. If data was not normal after an outlier analysis, a Kruskal-Wallis test was used to evaluate significant differences between data. If data was normal, equal variance assumption was tested between samples with a Browne-Forsythe test. If assumption was not met, a t-test with a Welch correction was used between two samples being compared. A t-test was used for all analysis between only two samples. If all assumptions of normality and equal variance were met and the power was above 0.8, then an ANOVA with a Tukey post-hoc test was used to determine significance. If the power was below 0.8 for any sample set, then the data was deemed inconclusive. The number of samples used for each group was based on previous studies utilizing similar collagen scaffolds (44,45). compressive testing (n=8), pore size (n=3), metabolic activity (n=6), cell number (n=5), western blot (n=3), gene expression (n=5), and micro-CT and histology (n=3). Error bars are represented as mean ± standard deviation.

## 3. Results

### 3.1. Addition of amniotic membrane matrix alters scaffold pore size and mechanical properties

The mineralized collagen scaffolds containing amniotic membrane matrix displayed qualitatively similar open-pore microstructures as seen in conventional mineralized collagen scaffolds (**Fig. 3B, C**). However, quantitative analysis of pore size revealed mineralized collagen-amnion scaffolds contain significantly (p < 0.05) smaller pore size (112 μm vs. 168 μm) compared to mineralized collagen scaffolds (**Fig. 3D, E**). Addition of amniotic membrane matrix also altered mechanical performance of the scaffold under compressive loading, with a notably significant (p< 0.05) increase in Young’s Modulus as well as values of collapse stress (σ*) and collapse strain (ε*) associated with the transition between linear elastic and collapse plateau regimes of low-density open-cell foam materials (**Fig. 4**).

**Figure 3.**
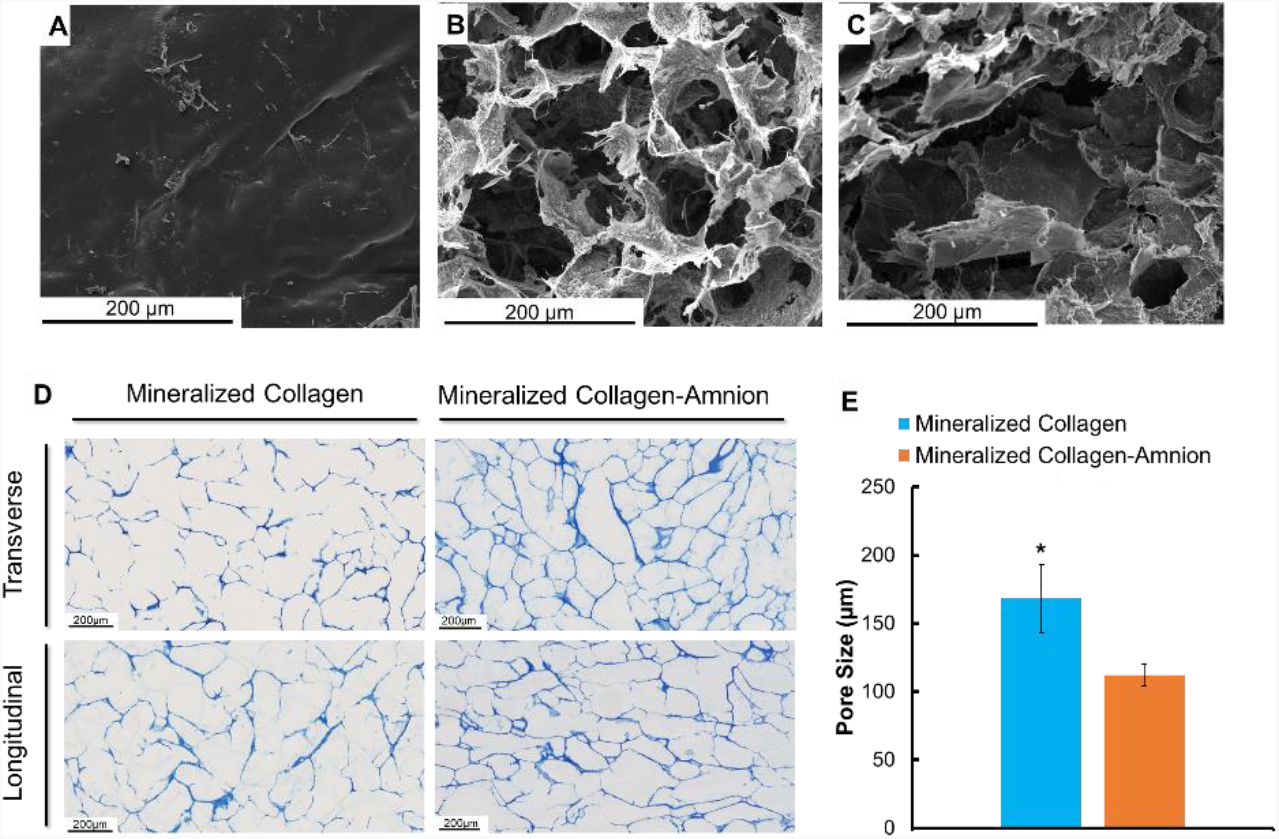
SEM imaging and pore size analysis of mineralized collagen and mineralized collagen-amnion scaffolds. (A) SEM image of the amniotic membrane. (B) SEM image of a mineralized collagen scaffold. (C) SEM image of a mineralized collagen-amnion scaffold. (D) Representative images of mineralized collagen and mineralized collagen-amnion scaffolds sectioned transversely and longitudinally and stained with aniline blue to determine pore size. (E) Average pore size of mineralized collagen and mineralized collagen-amnion scaffolds. * indicates the mineralized collagen scaffold has a significantly (p < 0.05) higher average pore size than the mineralized collagen-amnion scaffold. Data expressed as mean ± standard deviation (n=3).

**Figure 4.**
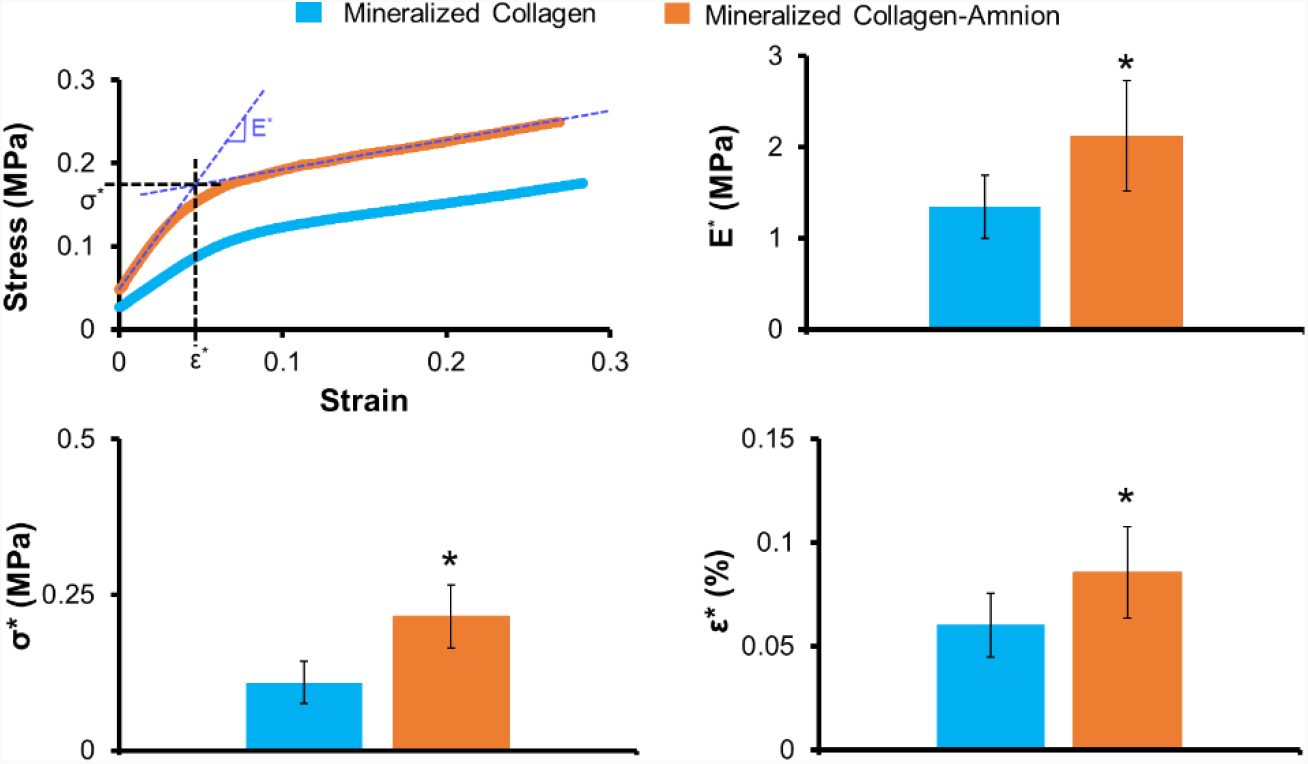
Mechanical compression testing of mineralized collagen and mineralized collagen-amnion scaffolds. Elastic Modulus is represented by E*, collapsed strain is represented by ε*, and collapsed stress is indicated by 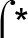. * indicates significantly (p < 0.05) greater average value of the mineralized collagen-amnion scaffold than the mineralized collagen scaffold. Data expressed as mean ± standard deviation (n=8).

### 3.2. The addition of the amniotic membrane to mineralized collagen scaffolds decreased the cell viability in scaffolds

Metabolic activity and cell number of mineralized collagen and mineralized collagen-amnion scaffolds in normal growth media were measured over the course of 28 days. The metabolic activity of pACS within the mineralized collagen-amnion scaffold showed a significant increase over the course of the 28-day experiment compared to the initial seeding density. However, pASC within the mineralized collagen scaffolds displayed a significantly (p < 0.05) greater cell metabolic activity across all days (**Fig. 5A**). Similarly, overall number of pASC increased in the mineralized collagen scaffolds over the course of 28 days, but the overall number of pASC within the mineralized collagen-amnion scaffolds remained constant over the course of the experiment (**Fig. 5A**).

**Figure 5.**
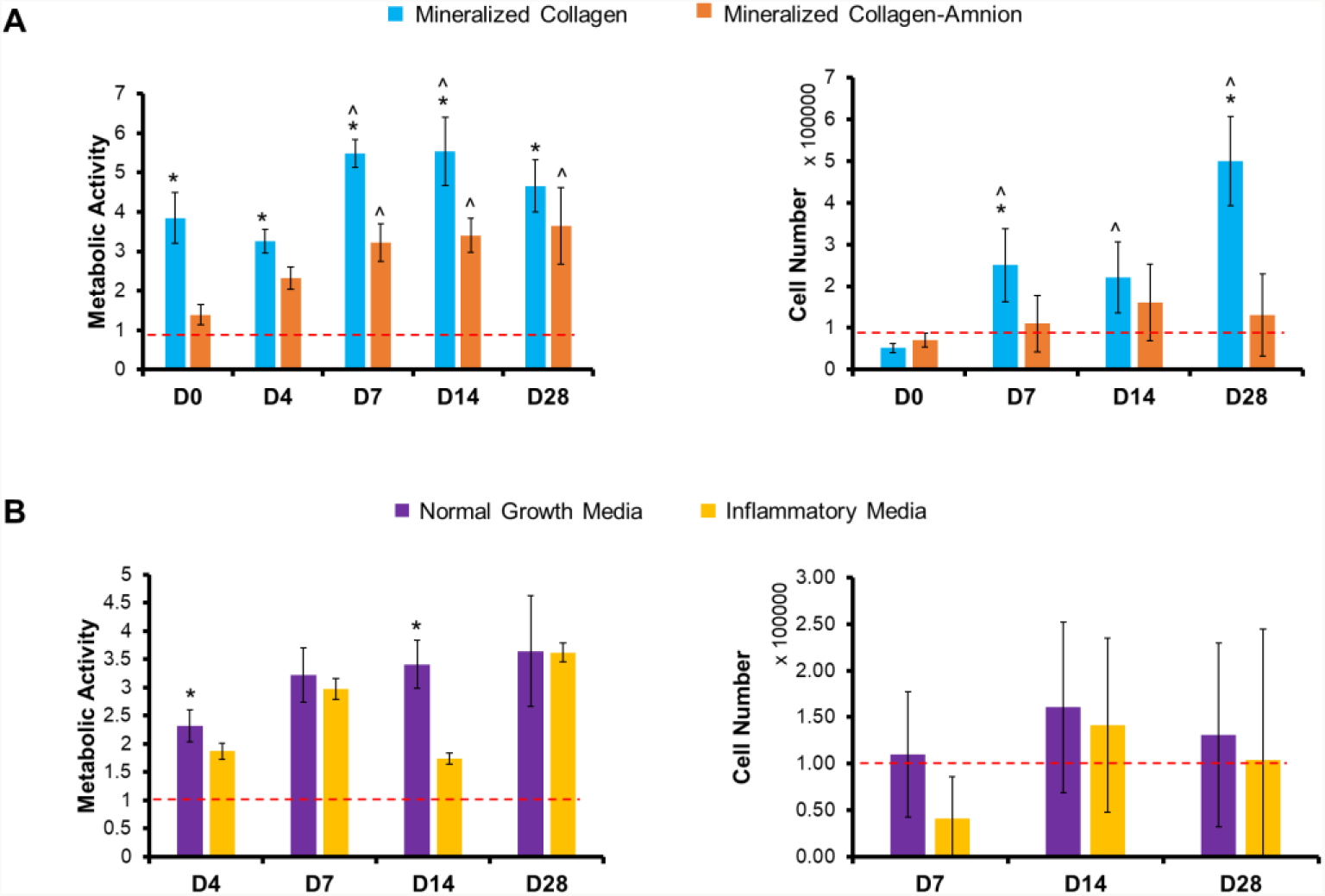
Cell viability of mineralized collagen and mineralized collagen-amnion scaffolds in normal growth media and inflammatory media. Metabolic activity was measured by an Alamar Blue assay, with a value of 1 representing the metabolic activity of 100,000 cells seeded on the scaffolds at the start of the experiment. Cell number was measured by a Hoechst DNA assay and 100,000 cells were initially seeded on the scaffolds. (A) Metabolic activity and cell number of mineralized collagen scaffolds compared to mineralized collagen-amnion scaffolds in normal growth media. (B) Metabolic activity and cell number of mineralized collagen-amnion scaffolds in normal growth media and inflammatory media. * indicates the metabolic activity or cell number of the mineralized collagen scaffolds is significantly (p < 0.05) greater than the mineralized collagen-amnion scaffolds on the same day. ^ indicates the metabolic activity or cell number of one scaffold type was significantly (p < 0.05) greater than the same scaffold type at day 0. Data expressed as mean ± standard deviation (Alamar Blue: n=6, Hoechst: n=5).

### 3.3. Addition of inflammatory challenge does not impact long-term cell viability within mineralized collagen-amnion scaffolds

We subsequently examined whether inclusion of amniotic membrane matrix within the mineralized collagen scaffold reduced the deleterious effect of 1 ng/mL soluble IL-1β upon pASC bioactivity, monitoring pASC metabolic activity and cell number over the course of 28 days. While some significant differences in metabolic activity were observed between inflammatory challenge and conventional media at individual timepoints, overall, we observed no deleterious effect of IL-1β on pASC metabolic activity or overall number (no significant differences between the groups at any of the timepoint) within mineralized collagen-amnion scaffolds throughout the entire 28 day experiment (**Fig. 5.B**).

### 3.4. Protein activity was unchanged dependent on scaffold type and media condition

We subsequently examined activity of AKT, p38, and ERK1/2 in mineralized collagen scaffolds in normal growth media as well as mineralized collagen-amnion scaffolds in response to growth or inflammatory media. These proteins were selected for their involvement in the bone formation and mineralization process and prior identification of their activation in conventional mineralized collagen scaffolds. We observed no significant (p < 0.05) differences between the two scaffold types in normal growth medium for all proteins examined (**Supp. Fig. 2**). P38 activity was expressed at near 100% at the early timepoints in both scaffold types and ERK1/2 and AKT were expressed at near 100% at later timepoints. Additionally, we observed no significant (p < 0.05) differences in protein activity of mineralized collagen-amnion scaffolds in response to inflammatory challenge (**Supp. Fig. 2**).

### 3.5. Mineralized collagen scaffolds have increased osteogenic transcription factor *RUNX2* compared to mineralized collagen-amnion scaffolds, but no differences in expression of other osteogenic genes

We examined baseline shifts in gene expression patterns for a library of osteogenic genes (*RUNX2, BGLAP, COL1A2*, *BMP2*) in mineralized collagen vs. mineralized collagen-amnion scaffolds. *RUNX2* was significantly upregulated in mineralized collagen scaffolds and was even more upregulated than in mineralized collagen-amnion scaffolds (**Supp. Fig. 3**). However, there were no significant (p < 0.05) differences in *BGLAP, COL1A2*, or *BMP2* expression between the two scaffolds.

### 3.6. Osteogenic genes are increased in mineralized collagen-amnion scaffolds in inflammatory media

We subsequently examined whether inclusion of inflammatory challenge altered osteogenic gene expression profiles in the mineralized collagen-amnion scaffolds. Interestingly, *RUNX2* and *COL1A2* were significantly (p < 0.05) upregulated in the presence of inflammatory medium compared to normal growth media at days 14 and 28 (**Fig. 6**). *BGLAP* and *BMP2* showed 15-fold and 2-fold, respective increases in expression at day 28 in inflammatory medium after showing little to no upregulation in normal growth media.

**Figure 6.**
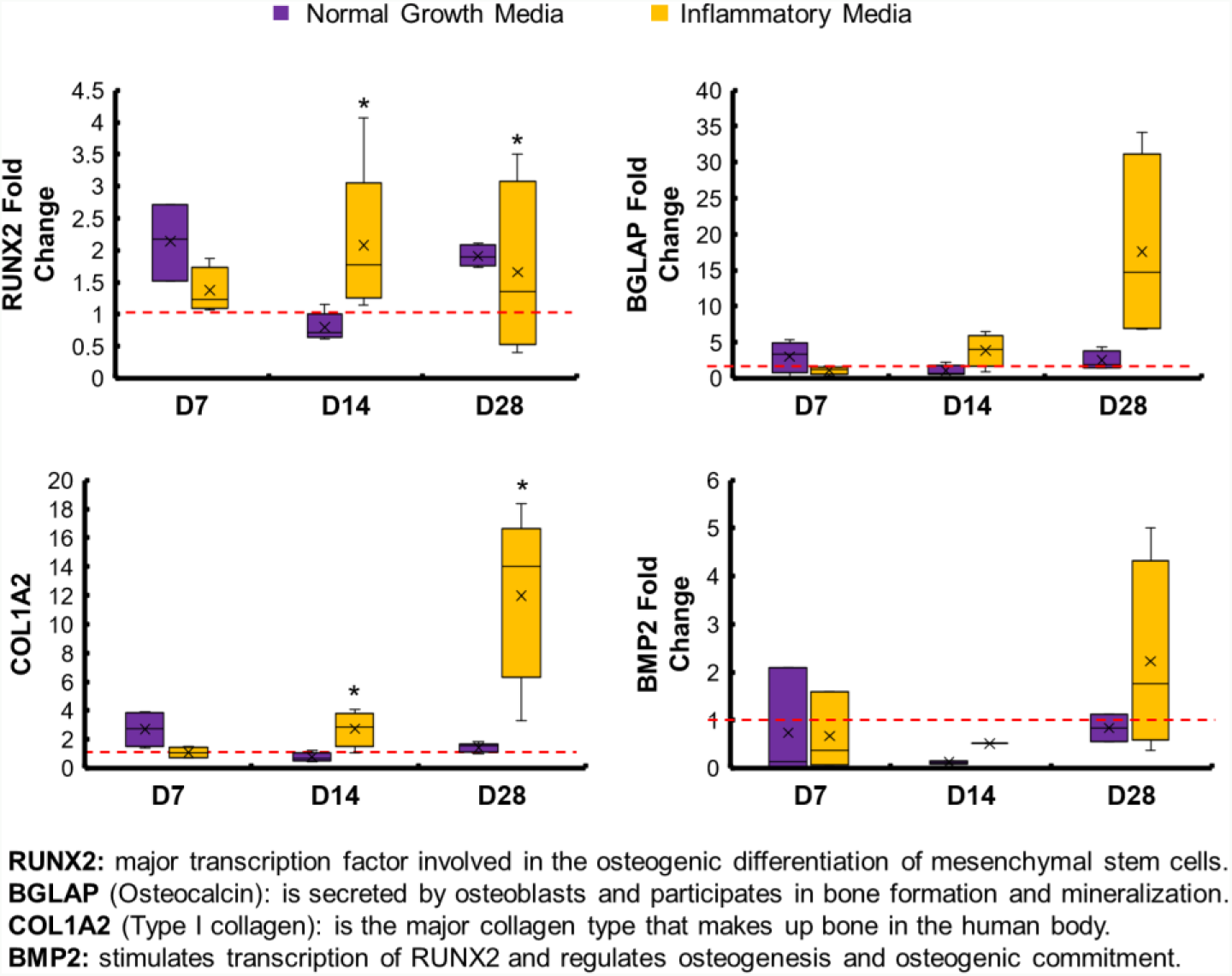
Osteogenic gene expression of mineralized collagen-amnion scaffolds in normal growth media and inflammatory media. Gene expression was evaluated by RT-PCR and normalized to the expression of cells before seeding on scaffolds. * indicates the mineralized collagen scaffold was significantly (p < 0.05) greater than the mineralized collagen-amnion scaffold on the same day. Data expressed as mean ± standard deviation (n=5).

### 3.7. Mineral remodeling was enhanced in mineralized collagen amnion scaffolds in response to inflammatory challenge

Finally, we examined the influence of amniotic membrane matrix and inflammatory challenge on mineralization and remodeling via Micro-CT and immunohistochemical analysis. In all cases, results after 28 days in culture were compared to unseeded (day -1) control scaffolds. After 28 days in normal growth media, we observed no significant (p < 0.05) difference in overall changes in mineral volume fill via μCT between mineralized collagen and mineralized collagen-amnion scaffolds (**Fig. 7A**). As a result, we performed immunohistochemical analyses of Eosin, Alizarin Red S, and Von Kossa to define mineral content, cells, and pore architecture of the scaffolds. Von Kossa staining revealed dark mineral staining and with many visible cells around the stained collagen in both mineralized collagen and collagen-amnion scaffolds (**Supp. Fig. 4A**). Hematoxylin and Eosin revealed both variants retained an open porous network after 28 days, while Alizarin Red S stains demonstrated darker red staining in the mineralized collagen samples than the mineralized collagen-amnion samples suggesting more calcium present (**Supp. Fig. 4A**). Interestingly, we did observe a significant (p < 0.05) increase in mineral volume fill in mineralized collagen-amnion scaffolds in response to inflammatory challenge (**Fig. 7B**). Examining the explicit effect of inflammatory cytokines on pASC activity, we observed consistent Von Kossa and fainter Alizarin Red S staining in scaffolds exposed to inflammatory challenge (**Supp. Fig. 4B**). Lastly, we observed OPN positive cells in both mineralized collagen and mineralized collagen-amnion scaffolds, concentrated mostly towards the periphery of the scaffold (**Supp. Fig. 5**). The inclusion of inflammatory challenge did not affect the presence of OPN positive cells on mineralized collagen-amnion scaffolds and OPN-positive cells were present throughout these scaffolds.

**Figure 7.**
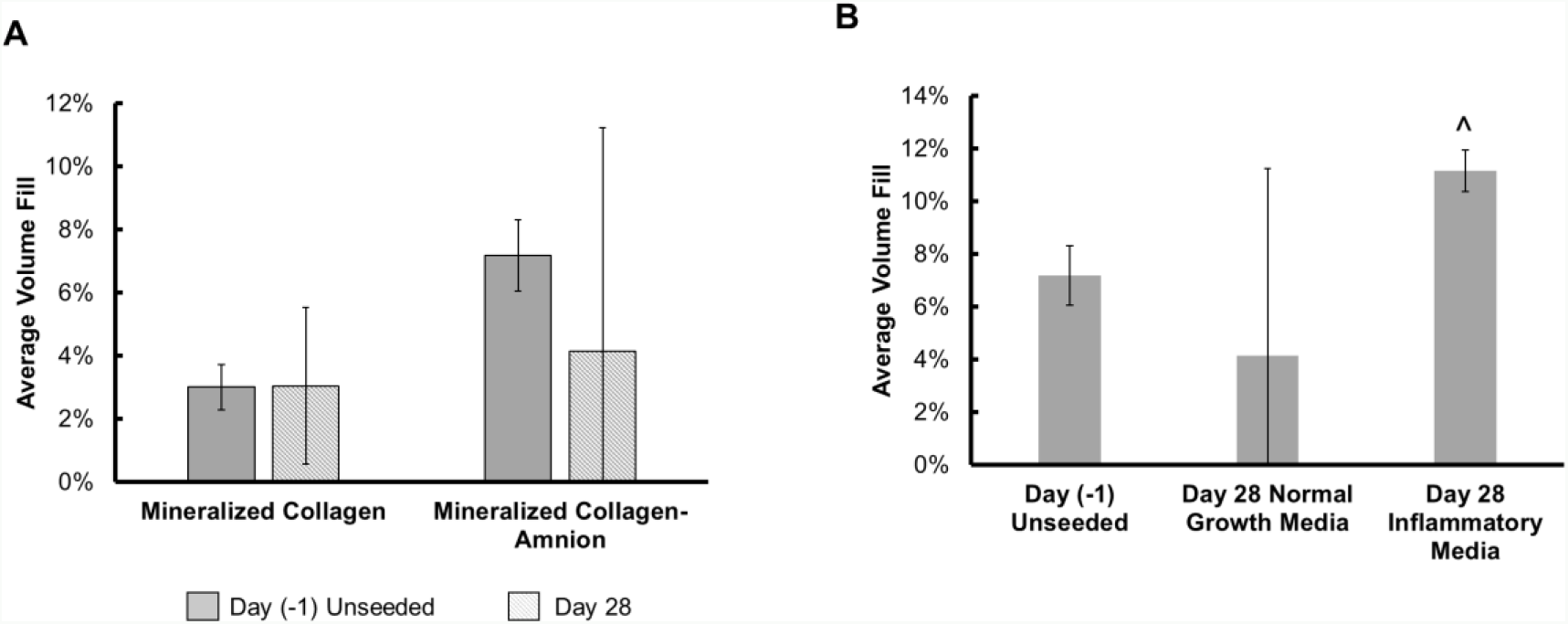
Mineral formation in mineralized collagen and mineralized collagen-amnion scaffolds in normal growth media. Average mineral fill was quantified by ImageJ processing of micro-CT stacks of mineralized collagen and mineralized collagen-amnion scaffolds. (A) Mineral formation of mineralized collagen scaffolds and mineralized collagen-amnion scaffolds in normal growth media without cells at day (−1) and with cells at day 28. No significance was found between all samples. (B) Mineralized collagen-amnion scaffolds in normal growth media and inflammatory media. ^ indicates the day 28 inflammatory media group was significantly (p < 0.05) greater than the day (−1) unseeded group. Data expressed as mean ± standard deviation (n=3).

## 4. Discussion

In this study, we describe the fabrication of a novel mineralized collagen scaffold variant to include human amniotic membrane matrix derived from placentas. We report the influence of amniotic membrane matrix inclusion on cell osteogenesis and activity *in vitro* of these scaffolds in response to inflammatory cytokine challenge. Regenerative solutions for complex CMF defects will likely require the use of biomaterials that promote osteogenesis and modulate the inflammatory response to accelerate healing. Herein, we address the challenge of inflammation by utilizing the amniotic membrane, which has been known to be anti-inflammatory (27–31). Modulating the inflammatory response of implants can increase the likelihood of success, by preventing fibrous encapsulation of implants which can occur due to persistent inflammation (5). We have recently developed a mineralized collagen scaffold that natively promotes osteogenic differentiation and osteogenesis of various cell types in the absence of osteogenic supplements and osteogenic media (8,18). These scaffolds promote activation of SMAD1/5/8, ERK1/2, AKT, p38 MAPK pathways, which ultimately lead to mineral formation both *in vivo* and *in vitro* (16,48,49). Although these scaffolds are successful in many sub-critical sized defects (26,48), larger defects still remain a challenge. These mineralized collagen scaffolds do not contain any immune-regulatory components, such that there still exists a risk of chronic inflammation. To address this challenge, we explored the addition of the amniotic membrane, an ECM containing anti-inflammatory properties. This was previously explored in our lab in combination with non-mineralized collagen scaffolds for tendon repair with promising outcomes (37,38). However, the use of non-mineralized collagen scaffolds in CMF bone defect repair does not provide necessary mineralization and hence, adaptations to the mineralized collagen scaffolds must be made. Here, we address this critical need of immune regulation by the successful addition of the amniotic membrane to mineralized collagen scaffolds.

We described the incorporation of the amniotic membrane (5:1 collagen:amnion content) into mineralized collagen scaffolds to create a mineralized collagen-amnion scaffold. We report the activity of porcine adipose derived stem cells in mineralized collagen-amnion scaffolds in response to basal media vs. inclusion of inflammatory challenge (1 ng/mL IL-1β (37)). We hypothesized the addition of the amniotic membrane matrix to mineralized collagen scaffolds would increase osteogenic activity despite inflammatory challenge. To test this hypothesis, we examined our scaffolds for amniotic membrane incorporation, *in vitro* mineral formation, and osteogenic gene and protein response. With the addition of the amniotic membrane, the mineralized collagen scaffolds still demonstrated an open porous network similar to mineralized collagen scaffolds. While mineralized collagen-amnion scaffold show reduced pore size, the pores of the mineralized collagen-amnion scaffolds remain in an appropriate range of cell infiltration for bone regeneration (100-350μm) (50). Scaffold mechanical properties were also affected, with the addition of the amnion leading to increase in all compressive properties (Young’s Modulus, collapse stress, collapse strain). Structural changes in the scaffolds could be attributed to the amount of amniotic membrane added. The amount added was based on previous designs of collagen-amnion scaffolds (37), using a 5:1 ratio of collagen:amnion, with the collagen content being much higher in mineralized collagen scaffolds than non-mineralized collagen scaffolds and thus requiring more amnion to be used. Lastly, results from pore size and mechanical analysis as well as SEM imaging (**Fig. 3**) suggest that the amniotic membrane matrix was distributed throughout the scaffold microstructure.

We subsequently profiled pASC bioactivity and osteogenic potential in mineralized collagen scaffolds as a function of amniotic membrane matrix. Cell viability was negatively affected by the addition of the amnion (**Fig. 5. A**); however, scaffolds containing the amniotic membrane showed overall increase in metabolic activity over 28 days, indicating that although the cells were more viable on scaffolds lacking the amniotic membrane, the presence of amniotic membrane matrix was not cytotoxic. Osteogenic gene expression suggested pASC retained their osteogenic potential in the mineralized collagen-amnion scaffolds. *RUNX2* was significantly (p < 0.05) more upregulated at day 0, 14, and 28 compared to scaffolds containing the amniotic membrane, with a near 4-fold change at day 28. However, all other osteogenic genes examined had few differences between the two scaffold types (**Supp. Fig. 3**). *BGLAP* is secreted by mature osteoblasts and can be used as a marker for bone mineralization, and both scaffold types were upregulated throughout the study. *COL1A2*, a marker for type I collagen, also had no differences between the two groups, and most days besides day 14 were upregulated. *BMP2*, which stimulates the transcription of *RUNX2* (51,52), had no differences between groups but was downregulated at most days. In addition, there were no significant (p < 0.05) differences in expression of osteogenic signaling proteins, AKT, p38, and ERK1/2 (**Supp. Fig. 2**). Micro-CT revealed no differences in mineral formation at the end of the study between the two groups, but highlighted more variable mineral formation in scaffolds containing the amniotic membrane. Mineral staining using Von Kossa and Alizarin Red S demonstrated the presence of calcium in both scaffold types and an open porous structure (**Supp. Fig. 4**). Osteopontin is expressed in various bone cells and is important for mineralization and formation, as well as bone remodeling (53,54). Osteopontin-positive cells were visible throughout both scaffold types, as indicated by black arrows (**Supp. Fig. 5**). In sum, pASC seeded on scaffolds containing the amniotic membrane have similar late-stage mineralization compared to those without, but with less osteogenic differentiation, suggesting mineralized collagen-amnion scaffolds have potential as an osteogenic implant.

More importantly, we examined whether the presence of inflammatory conditions reduced the osteogenic potential of the mineralized collagen-amnion scaffolds. While the inclusion of an inflammatory challenge may have temporarily reduced pASC metabolic activity (days 1-4), over extended culture (day 28) the inclusion of amniotic membrane abrogated any deleterious effects of persistent IL1-β challenge. Interestingly, osteogenic genes *RUNX2* and *COL1A2* were both significantly upregulated in amnion scaffolds in response to inflammatory challenge (**Fig. 6**). Together, gene and protein expression indicated that cells experiencing inflammatory conditions on the amniotic membrane promoted more osteogenic differentiation. Micro-CT and mineral staining via immunohistochemistry further verified this, where inflammatory-challenged mineralized collagen-amnion scaffolds showed significant increases in mineral deposition. Overall, the presence of inflammatory medium positively influenced osteogenesis responses, without hindering the osteogenic potential of the mineralized collagen amnion scaffolds.

We report inclusion of amnion membrane derived matrix can be included within a model collagen scaffold to enhance osteogenic potential in response to inflammatory cytokines. Inflammation is an essential part of healing CMF defects, and a material that could promote osteogenesis and healing in these conditions may improve regenerative potential. The amniotic membrane has been previously characterized to have glycosaminoglycans (0.22 wt %) and high collagen content (40.8 wt %) (37). Future work will further characterize this membrane to determine types of collagen present and biomolecule release, as well as using the amniotic membrane as the main supply of collagen in new scaffold designs. Scaffold mechanics were affected by the amniotic membrane, thus future work will involve incorporating finer particles of this membrane in order to maintain the same pore size and mechanical properties. Interestingly, work by Go et al., suggested extracts from chorion vs. amnion membrane may more efficiently promote osteogenic responses (32,36). As we found limited osteogenic differences between mineralized collagen and mineralized collagen-amnion scaffolds, ongoing work is exploring the use of amniotic membrane vs. chorionic membrane matrix in this scaffold system. Lastly, this work focused on the deleterious effect of inflammatory challenge directly on an osteoprogenitor population. However, osteoprogenitors are only one element of a more complicated *in vitro* response that includes osteoclasts and members of the immune system, notably macrophages. We have recently shown the mineralized collagen scaffold can both directly, and indirectly via secretome generated by osteprogenitors, transiently inhibit osteoclast activity (55,56). Ongoing studies are examining the influence of inclusion of amniotic membrane matrix within the scaffold on the polarization kinetics of macrophages to enhance the capacity of the mineralized scaffold to accelerate CMF bone regeneration.

## 5. Conclusions

We report the incorporation of amniotic membrane matrix into a mineralized collagen scaffolds under development for CMF bone regeneration. The addition of the amniotic membrane increased compressive properties and decreased pore size. Cell viability and osteogenic differentiation were greater in mineralized collagen scaffolds without the amniotic membrane, but mineral formation volume was the same at the end of the study. Amniotic membrane-containing scaffolds also demonstrated improved osteogenesis and mineral formation in response to inflammatory challenge. The addition of the amniotic membrane to mineralized collagen scaffolds indicates a potential to improve osteogenic repair of CMF defects even under inflammatory conditions.

## Acknowledgements

The authors would like to acknowledge the Beckman Institute for Advanced Science and Technology at the University of Illinois at Urbana-Champaign, especially the Imaging Technology Group and members Leilei Yin (Micro-CT), Scott Robinson and Cate Wallace (ESEM). The authors would also like to acknowledge the Carl R. Woese Institute for Genomic Biology for the use of their core instruments, including Kingsley Boateng for help with histology preparation. The authors would like to acknowledge the Roy J. Carver Biotechnology Center and assistance with RT-PCR from Tatsiana Akraiko as well as Carle Medical Hospital for providing placental tissues. In addition, the authors would like to thank Rebecca Hortensius for assistance with isolation of human placentas, and Samantha Zambuto and Aleczandria Tiffany for assistance with manuscript preparation. This work was supported by the Office of the Assistant Secretary of Defense for Health Affairs Broad Agency Announcement for Extramural Medical Research through the Award No. W81XWH-16-1-0566. Opinions, interpretations, conclusions and recommendations are those of the authors and are not necessarily endorsed by the Department of Defense. Research reported in this publication was also supported by the National Institute of Dental and Craniofacial Research of the National Institutes of Health under Award Number R21 DE026582. The content is solely the responsibility of the authors and does not necessarily represent the official views of the NIH. We are grateful for the funding for this study provided by the NSF Graduate Research Fellowship DGE-1144245 (MD).

