## Supplemental Info for "Mineralized collagen scaffolds fabricated with amniotic membrane matrix increase osteogenesis under inflammatory conditions"

[1] Dept. of Materials Science and Engineering
[2] Dept. of Bioengineering
[3] Dept. Chemical and Biomolecular Engineering
[4] Carl R. Woese Institute for Genomic Biology
[5] Dept. of Animal Sciences

University of Illinois at Urbana-Champaign
Urbana, IL 61801

**Supplementary Figures**

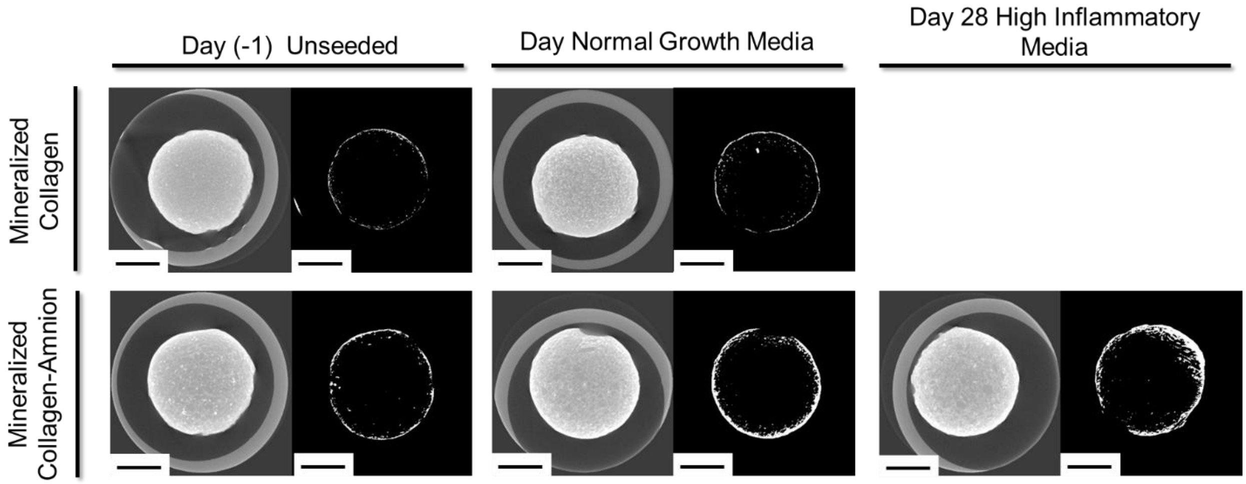


**Supplementary Figure 1. Representative Micro-CT images and ImageJ renditions of mineralized collagen and mineralized collagen-amnion scaffolds.** Raw micro-CT images presented on the left of each column and ImageJ intensified images presented on the right. Scale bar represents 2.5 mm.

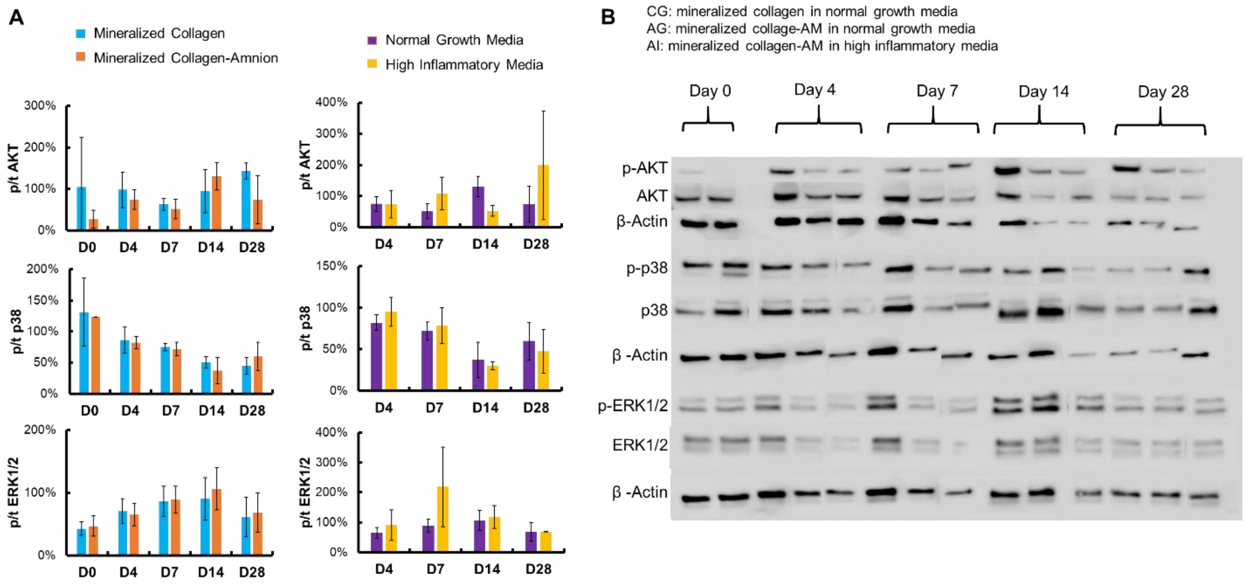


**Supplementary Figure 2. Protein activity of mineralized collagen scaffolds compared to mineralized collagen-amnion scaffolds in normal and high inflammatory media.** Osteogenic protein activity was quantified with Western Blots. (A) Phosphorylated over total protein activity in scaffolds. No significance ($p < 0.05$) was observed between any groups. Data expressed as mean ± standard deviation (n= 3). (B) Representative western blot images with β-actin as a control.

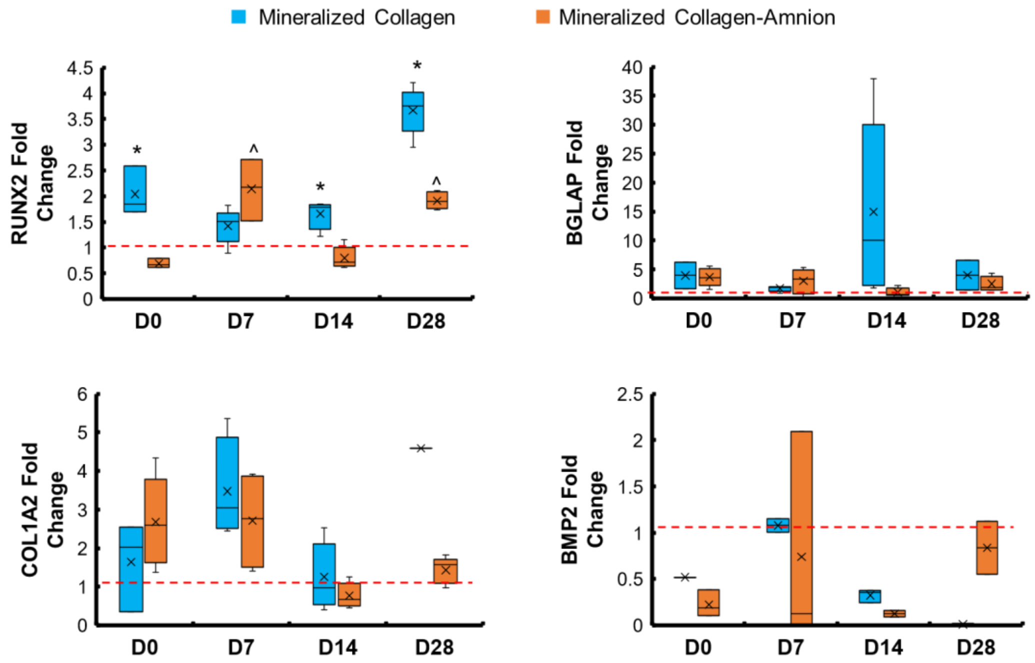


**Supplementary Figure 3. Osteogenic gene expression of mineralized collagen and mineralized collagen-amnion scaffolds in normal growth media.** Gene expression was evaluated by RT-PCR and normalized to the expression of cells before seeding on scaffolds. Below the graphs are brief statements about each gene of interest. * indicates the mineralized collagen scaffold was significantly ($p < 0.05$) greater than the mineralized collagen-amnion scaffold on the same day. ^ indicates one scaffold type was significantly ($p < 0.05$) greater than the same type compared to day 0. Data expressed as mean ± standard deviation (n=5).

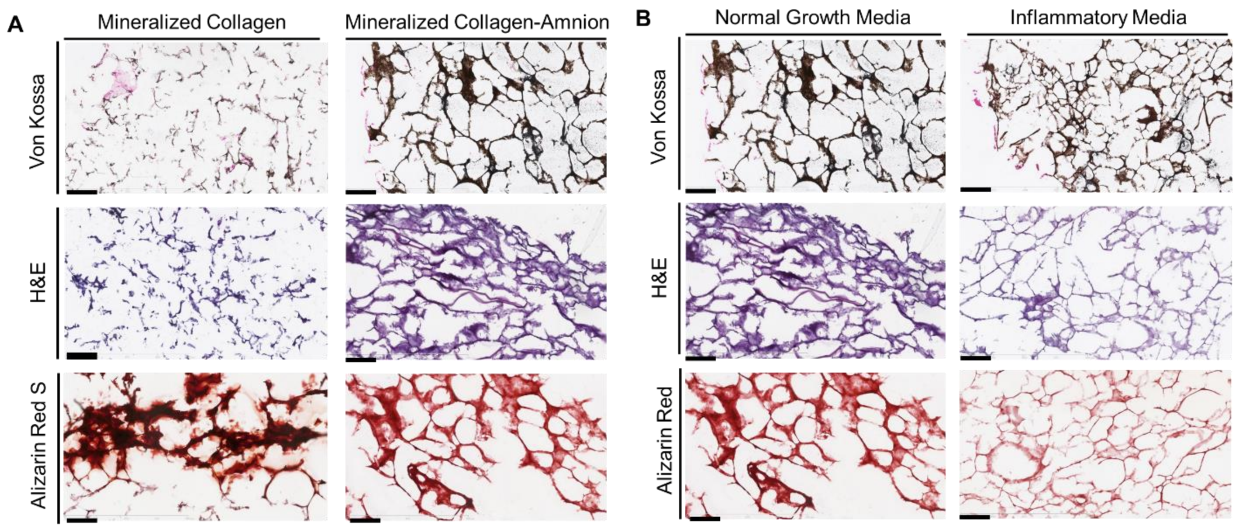


**Supplementary Figure 4. Histological staining of mineralized collagen-amnion scaffolds in normal growth media and inflammatory media after 28 days.** (A) Mineralized collagen and mineralized collagen-amnion scaffolds in normal growth media. (B) Mineralized collagen-amnion scaffolds in normal growth media and inflammatory media. Scale bar represents 200µm.

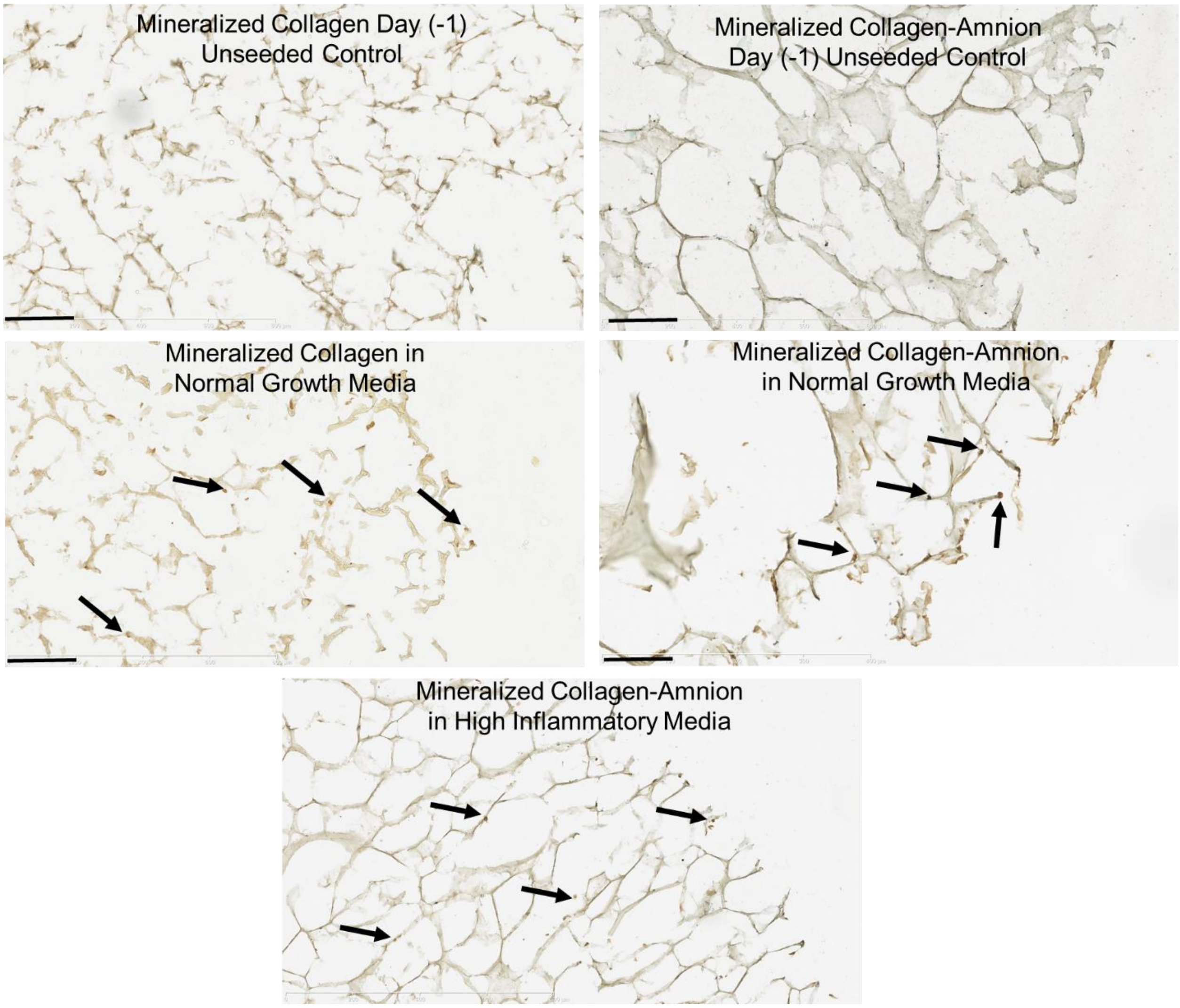


**Supplementary Figure 5. Osteopontin-stained mineralized collagen and mineralized collagen-amnion scaffolds.** Scaffolds were stained using immunohistochemical procedures. Black arrows represent OPN positive cells. Scale bars represent 200 µm.
